# A “grappling hook” interaction balances self-assembly and chaperone activity of Nucleophosmin 1

**DOI:** 10.1101/2022.09.29.510028

**Authors:** Mihkel Saluri, Axel Leppert, Genis Valentin Gese, Cagla Sahin, Dilraj Lama, Margit Kaldmäe, Gefei Chen, Arne Elofsson, Timothy M. Allison, Marie Arsenian-Henriksson, Jan Johansson, David P. Lane, B. Martin Hällberg, Michael Landreh

## Abstract

How the self-assembly of partially disordered proteins generates functional compartments in the cytoplasm and particularly in the nucleus is poorly understood. Nucleophosmin 1 (NPM1) is an abundant nucleolar protein that forms large oligomers which provide the scaffold for ribosome assembly but also prevent protein aggregation as part of the cellular stress response. Examining the relationship between the self-assembly and chaperone activity of NPM1, we find that oligomerization of full-length NPM1 modulates its ability to retard amyloid formation *in vitro*. Machine learning and cryo-electron microscopy reveal fuzzy interactions between the disordered region and the C-terminal nucleotide-binding domain that cross-link NPM1 pentamers into oligomers. Ribosomal peptides mediate in a tighter association within the oligomers, reducing their capacity to prevent amyloid formation. We conclude that NPM1 uses a “grappling hook” interaction to form a network-like structure whose chaperone activity is tuned by basic proteins, suggesting a regulatory mechanism for the nucleolar stress response.

## Introduction

The nucleolus, the site of ribosome biogenesis in the nucleus, is a membraneless compartment that responds to changes in cellular growth rate, metabolic activity, and stress ^1^. Its dynamic nature allows a constant exchange of proteins and nucleotides with the surrounding nucleus and the cytoplasm. For example, the nucleolus sequesters p14^ARF^ and its binding partner human double minute 2 homolog (HDM2), which otherwise ubiquitinylates the tumor suppressor p53 to allow cell cycle progression ^2^. Its ability to recruit and release proteins has been attributed to the fact that nucleolar assembly is driven by liquid-liquid phase separation (LLPS) of a highly enriched subset of nucleolar proteins ^3^. By exhibiting different phase-separating properties, these proteins account for the co-existence of nucleolar regions with distinct functions ^4^. Nucleophosmin 1 (NPM1, also known as B23) is the main component of the outermost nucleolar phase, the granular component, where ribosomes are assembled. Mutations in NPM1 are commonly associated with acute myeloid leukemia (AML) ^5^. It is the main nucleolar interaction partner for the p14^ARF^ tumor suppressor and the c-MYC oncoprotein, and knockdown experiments have identified NPM1 as a promising target for cancer therapy ^6,7^.

NPM1 has a modular structure with folded N-terminal and C-terminal domains (NTD, residues 1-120, and CTD, residues 240-294) linked by an intrinsically disordered region (IDR, residues 120-240) (Figure 1a). The N-terminal domain adopts a β-sheet sandwich fold and assembles into pentamers that can form end-to-end decamers ^8^. The NTD and IDR contain three highly acidic poly-D/E stretches (A1, A2, and A3), as well as two regions with predominantly basic residues (B1 and B2) ^9^. The CTD is a nucleotide-binding domain composed of thee α-helices and is the site of AML-related mutations, which result in removal of a nucleolar localization signal ^10^. Full-length (FL) NPM1 self-assembles into large oligomers *in vitro* ^9,11^ and in cancer cells ^12^. The addition of polyanionic or polycationic molecules, such as basic peptides or RNA, induce LLPS through heterotypic interactions with the opposing charges on FL NPM1 ^9^. The isolated NTD can also undergo LLPS by binding basic peptides via its acidic A1 region ^13^. In addition, oligomeric NPM1 engages in homotypic interactions through contacts between its charged regions, which can give rise to LLPS under crowding conditions and are modulated by ionic strength ^9,14^.

**Figure 1.**
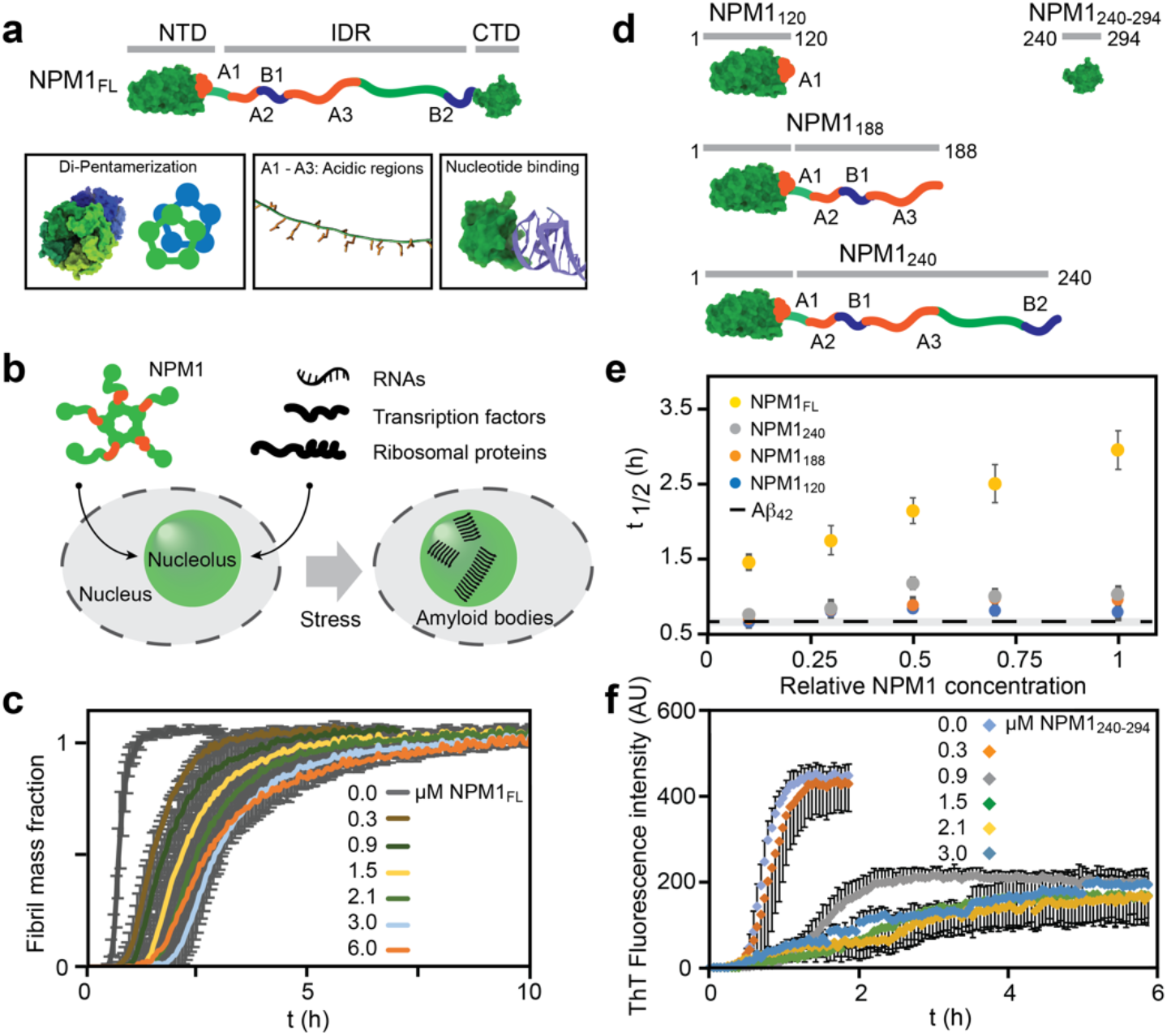
NPM1 delays Aβ_42_ aggregation. (a) NPM1 has a modular architecture, composed of a folded N-terminal pentamerization domain (NTD, residues 1-120), an intrinsically disordered region (IDR, residues 120-240) containing three acidic and two basic tracts (A1-3 and B1 and 2, respectively), and a C-terminal nucleotide-binding domain (CTD, residues 240-294). The NTD and the CTD are rendered based on PDB IDs 2P1B and 2VXD, respectively. (b) NPM1 pentamers associate with RNA, basic transcription factors, and ribosomal proteins to form the granular component of the nucleolus. Cellular stress induces the formation of nucleolar amyloid bodies. (c) The presence of increasing amounts of NPM1 shows that NPM1 delays the onset and the elongation of fibrillation, as judged by ThT fluorescence. Error bars indicate the standard deviation of n=4 experiments. (d) Truncated variants of NPM1 used in this study: NPM1_120_ encompasses the NTD. NPM1_188_ the NTD and the acidic regions of the IDR, NPM1_240_ the NTD and the entire IDR and NPM1_240-294_ only the CTD. (e) Fibrillation half-times (t _1/2_) of Aβ_42_ in the presence of 0 - 3 µM of NPM1 variants shows that only FL NPM1, but not truncated variants, affects fibrillation. The t _1/2_ of Aβ_42_ alone is shown as a dashed line. Error bars indicate the standard deviation of n=4 experiments. (f) ThT fluorescence curves of Aβ_42_ in the presence of 0 – 3 µM NPM1_240-294_ show a dose-dependent delay in fibrillation and a decrease in fluorescence intensity. Error bars indicate the standard deviation of n=4 experiments.

Besides its role as a nucleolar scaffold, NPM1 has a chaperone activity *in vitro* and effectively prevents aggregation of denatured proteins ^15^. Strikingly, the nuclear proteome is enriched in disordered and aggregation-prone proteins ^16^. The nucleolus can sequester misfolded proteins during cellular stress and turn them over to HSP70 for refolding ^17^. It also stores protein aggregates in amyloid bodies that may function as “nucleolar detention centers” for potentially toxic aggregates ^18^. In cells, NPM1 colocalizes with misfolded proteins and amyloids, suggesting that it functions as a chaperone during cellular stress ^17^ (Figure 1b). Importantly, deletion of individual domains of NPM1 reduces its ability to counteract thermal denaturation of proteins ^11^. Together, these findings raise the possibility that the chaperone function of NPM1 in the nucleolus is related to its self-assembly properties.

## Results

### Full-length NPM1 and its CTD display differential chaperone activity towards amyloid formation

Previous studies have established the ability of FL NPM1 to protect globular proteins from thermal and chemical denaturation ^11,15^. However, several amyloidogenic proteins are targeted to the nucleus ^19^, and NPM1 associates with nucleolar amyloid aggregates *in vivo* ^17^. We, therefore, tested the ability of NPM1 to prevent the aggregation of amyloid-β_1-42_ (Aβ_42_), which has been found in the nuclei of neurons from Alzheimer’s Disease patients and is considered a model for amyloid formation ^20,21^. Briefly, we incubated 3 µM Aβ_42_ with 0 – 6 µM NPM1 in the presence of the amyloid-specific dye Thioflavin T (ThT) and monitored fibril formation through the increase in fluorescence intensity (Figure 1c). The addition of NPM1 delayed the half-time of fibrillation in a concentration-dependent manner, with the most pronounced effect at an NPM1 : Aβ_42_ ratio of 1:1 (Figure 1c). The end point fluorescence intensity was in all cases comparable to that of Aβ_42_ alone (Figure S1a). Interestingly, NPM1 does not appear to specifically affect only primary or secondary nucleation, or fibril elongation, but rather all of these processes. Together, these observations suggest that NPM1 efficiently retards the amyloid assembly processes. To identify which parts of the modular NPM1 structure are responsible for delaying Aβ_42_ aggregation, we designed truncated variants: NPM1_120_, which contains only the NTD with the A1 tract, NPM1_188_, which includes the NTD and the acidic tracts A1-3, NPM1_240_, which encompasses the NTD and the entire IDR, and NPM1_240-294_, which represents the isolated CTD (Figure 1d). We then tested the effects of all NPM1 variants on Aβ_42_ fibrillation under the same conditions as for FL NPM1 and determined the half-time of Aβ_42_ fibrillation (t_1/2_). Plotting the Aβ_42_ t_1/2_ in the presence of FL NPM1, NPM1_120_, NPM1_188_, and NPM1_240_ shows that full-length NPM1 affects Aβ_42_ aggregation in a dose-dependent manner, whereas the C-terminally truncated variants have only minor effects (Figure 1e). NPM_240-294_, *i*.*e*. the isolated CTD, a dose-dependent delay in ThT fluorescence at a concentration of 0.3 µM, and additionally resulted in a strong suppression of ThT fluorescence when the concentration was raised further (Figure 1f). Strikingly, none of the other NPM1 constructs resulted in a similar decrease in end-point fluorescence (Figure S1a). In agreement with the ThT data, electron microscopy showed no fibrils in the presence of the CTD at the reaction end-point.

(Figure S1b). Together, these observatons indicate that the ability of NPM1 to delay fibril formation is related to its CTD but appears to be reduced in the FL protein. To test whether NTD and CTD work synergistically to delay Aβ_42_ fibrillation, we incubated Aβ_42_ with 3 µM NPM1_240_ and 3 µM NPM1_240-294_. However, the combined effect of both NPM1 parts on fibril formation was far less pronounced than for FL NPM1 at the same concentration (Figure S1b). This finding shows that the chaperone activity of the CTD is reduced in the presence of the NTD, but the effect is less pronounced when the NTD is part of the same polypeptide than when it is added *in trans*. These observations imply that inter-domain interactions in the FL protein regulate its chaperone activity towards Aβ_42_.

### A covalent link between NTD and CTD is required for NPM1 self-assembly

To better understand the relationship between chaperone activity and inter-domain interactions, we turned to native mass spectrometry (MS). Here, intact protein complexes are transferred from the solution to the gas phase using soft electrospray ionization (nESI). Since non-covalent interactions can be preserved during mass measurements, we can obtain information about the oligomeric states of protein complexes in solution ^22,23^. In combination with collision-induced dissociation, where the intact complexes are subjected to high-energy collisions with an inert buffer gas inside the mass spectrometer, we can determine their composition. By comparing the collision voltages required to dissociate protein interactions, we can furthermore assess their relative stabilities ^24^ (Figure 2a).

**Figure 2.**
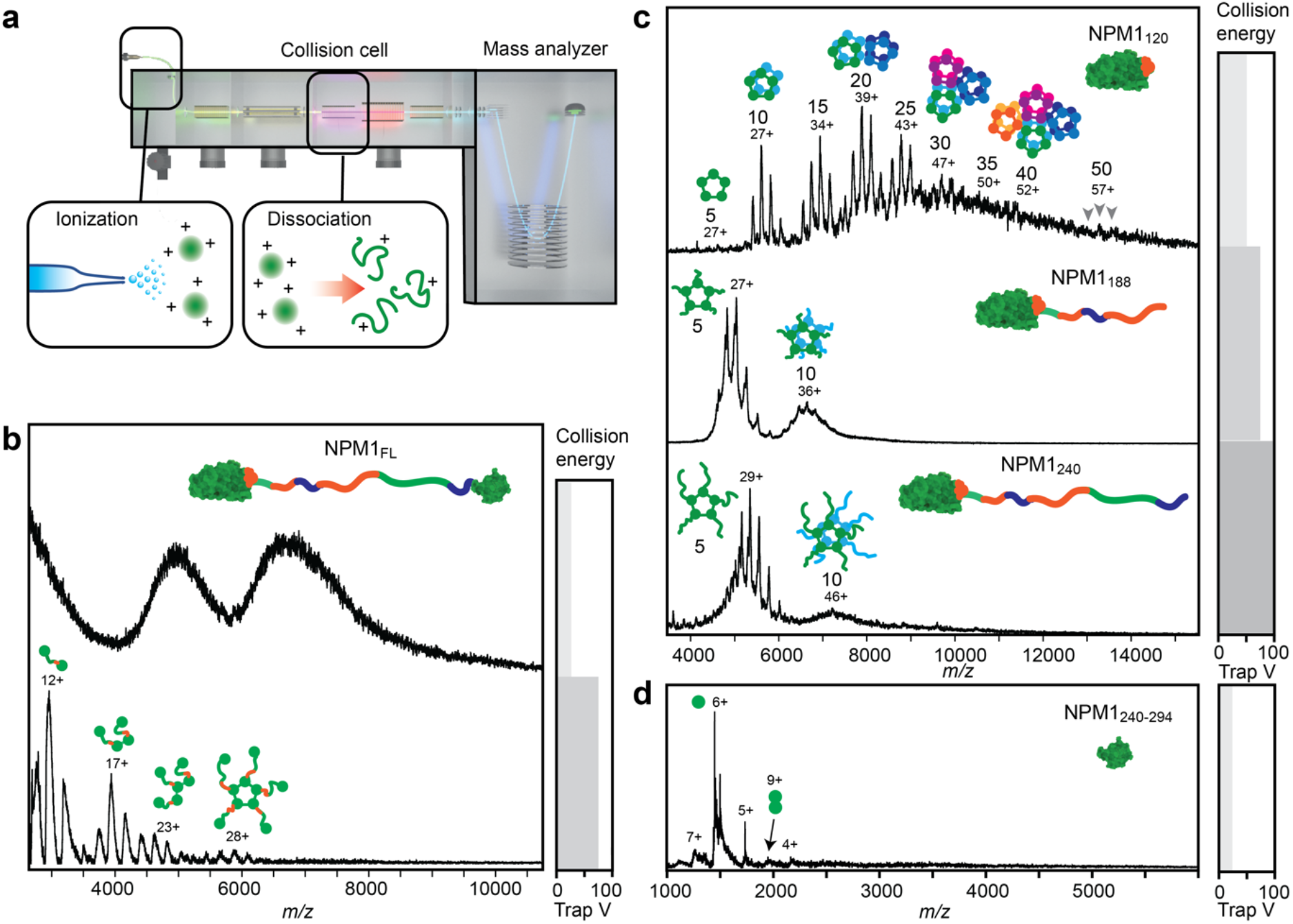
Native MS shows that higher oligomerization is specific for NPM1_FL_. (a) A schematic illustration of the native MS experiment, using collision-induced dissociation in the collision cell of the MS instrument. The illustration has been adapted from ^28^. (b) Native mass spectra of NPM1 at low collision energy show unresolved baseline humps, indicating large oligomers (top). Increasing the collision voltage releases NPM1 monomers, dimers, trimers, and pentamers (bottom). (c) Mass spectra of NPM1_120_ show a range of oligomeric states composed of multiples of five subunits as indicated. NPM1_188_ and NPM1_240_ form predominantly pentamers and minor decamer populations. (d) The isolated CTD (NPM1_240-294_) exists nearly exclusively as monomers. The respective collision voltage at which each spectrum was obtained is indicated on the right.

First, we subjected FL NPM1 to native MS analysis (Figure 2b). Under gentle MS conditions (collision voltage 25 V), we could only detect an unresolvable “hump” that is characteristic of large assemblies with no well-defined stoichiometry ^25^. When raising the collision voltage to 75 V, we obtained well-resolved peaks corresponding in mass to monomers, dimers, trimers, and pentamers of FL NPM1. We did not detect higher oligomeric states than pentamers and consider that the monomers, dimers, and trimers likely are dissociation products. These findings indicate that FL NPM1 pentamers assemble into large oligomers, in agreement with previous reports ^9,11,12^. Next, we analyzed oligomerization of the truncation variants. NPM_120_ yielded well-resolved spectra at a collision voltage of 50 V. Strikingly, we observed a broad range of oligomeric states, ranging from 5 to 50 subunits, but always in multiples of five, suggesting that the NTD pentamers can assemble into polymers (Figure 2b). The charge state distribution of a protein ion is dependent on its surface area ^26,27^, allowing structural information to be extracted. Plotting the average charge of each oligomeric state as a function of its molecular weight results in a correlation which closely follows the trend expected for globular proteins (Figure S2a). The crystal structure of NPM1_120_ shows side-by-side and end-to-end association of NTD pentamers in a crystal lattice via salt bridges (Figure S2b), which leads us to speculate that crystal-like interactions drive multimerization of the NTDs. MS analysis of NPM1_188_, which includes the IDR region with acidic tracts A2 and A3, required moderately higher collision voltages (75 V). The resulting mass spectra show pentamers and decamers, but not any higher oligomeric states, indicating that the IDR disrupts the self-association propensity of the isolated NTDs. Native MS analysis of NPM1_240_, which includes the entire IDR, yielded essentially the same oligomeric state as NPM1_188_, but with a slightly lower amount of decamers. Lastly, we also examined the isolated CTD, NPM1_240-294_. Native mass spectra show monomeric protein, with no sign of higher oligomerization besides traces of dimers (Figure 2d). Taken together, native MS reveals three types of NPM1 oligomerization, which are governed by the three domains: (1) the isolated NTD forms pentamers that self-assemble into ordered multimers; (2) the IDR disrupts these NTD multimers, yielding pentameric protein; (3) including the CTD induces the formation of large oligomers which can be dissociated into pentamers and pentamer fragments. We conclude that the formation of higher NPM1 oligomers requires full-length protein, suggesting direct interactions involving the CTD.

### NPM1 pentamers connect via their CTDs

The observations from MS suggest that the CTD is involved in higher oligomerization of NPM1. We therefore used fluorescence spectroscopy to test for interactions between the CTD and other regions of the protein. Importantly, the only two tryptophane residues in NPM1 are in the CTD, which allowed us to use intrinsic fluorescence to probe its interaction in solution. We found that the addition of quadruplex DNA, a high-affinity ligand for the CTD ^29^, quenches the intrinsic fluorescence of NPM1_240-294_, indicating binding (Figure 3a). We then tested whether the N-terminal region of NPM1 affected tryptophane fluorescence. Addition of NPM1_120_ resulted in a mild decrease in fluorescence intensity, whereas the addition of NPM1_240_ had a more pronounced effect. The presence of both DNA and NPM1_240_ gave an intermediate fluorescence reduction. Taken together, the change in tryptophane fluorescence suggests a direct association between NPM1_240-294_ and NPM1_240_, which is impacted by the presence of DNA.

**Figure 3.**
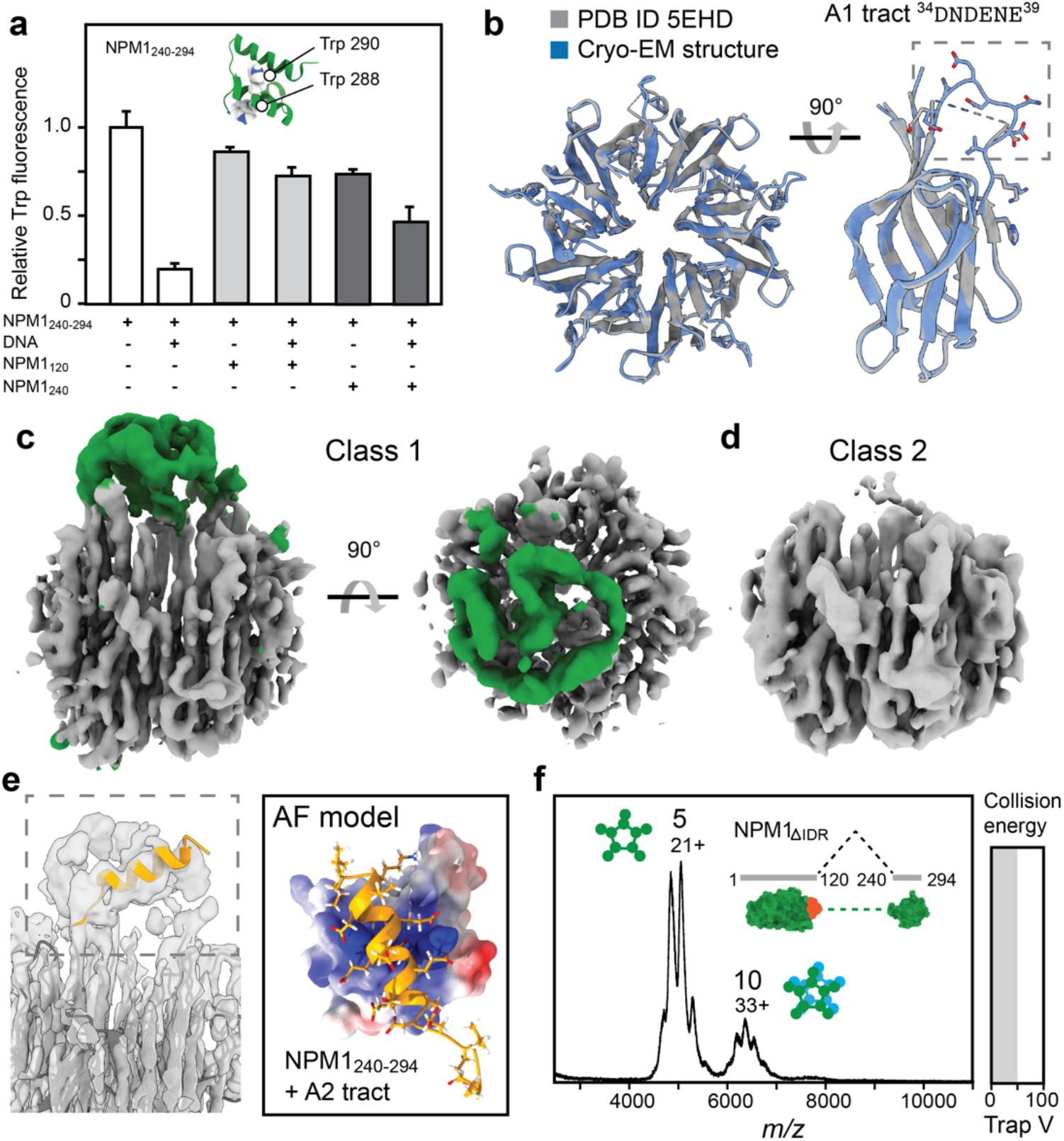
An NTD-CTD interaction in FL NPM1. (a) The fluorescence of tryptophane residues 288 and 290 in NPM_240-294_ is quenched by the addition of equimolar amounts of DNA or NPM_240_, and to a lesser extent by NPM_120_. In the presence of DNA and NPM_240_, intermediate quenching is observed. Error bars indicate the standard deviation of six experiments. (b) The cryo-EM reconstruction of the NTD (grey) shows virtually no deviations from the crystal structure (PDB ID 5EHD, blue), except for the A1 tract (residues 34-39), which could not be modeled based on the density map. (c) The cryo-EM density map for FL NPM1 reveals an additional asymmetric density (green) above the NTD pentamer (grey). (d) A second 3D class with fewer neighboring particles does not show the additional density. (e) The helical A2 tract (orange) appears as a diffuse density in the EM map. AF predicts a complex between the helical A2 tract (orange) and the basic CTD (rendered as electrostatic surface). (f) Native MS of the NPM_ΔIDR_ variant lacking the disordered region between residues 120-240 reveals pentamers and a small fraction of decamers, but no higher oligomers.

To better understand how the CTD associates with other parts of NPM1, we turned to cryo-electron microscopy (cryo-EM). Using physiological conditions (pH 7.5 and 150 mM NaCl), we obtained electron density maps that allowed a reconstruction of the NTD with a resolution of 2.6 Å (Figure S3, Table S1). Comparison to the crystal structure revealed a virtually identical fold, including sidechain orientations, except for the loop covering residues 34-39, which makes up the A1 tract. In the X-ray structure, it is stabilized via crystal contacts with the neighboring pentamer (Figure S2). In the cryo-EM density maps, the loop is too flexible for a confident reconstruction, suggesting that not only the A2 and A3 tracts, but even the A1 tract in the NTD is disordered (Figure 3b). Interestingly, we obtained two distinct 3D classes, one with an additional density above the acidic side of the pentamer, off-center from central cavity (Figure 3c). Strikingly, nearly all particles in this class had another NTD pentamer in proximity (Figure S4a). Particles in the second class, on the other hand, did not show the extra density, and had less nearby particles (Figure 3d, Figure S4b). The extra density is connected to the pentamer via the A2 tract of a single NTD and can therefore not be attributed to the disordered regions of each protomer (Figure S4c). We speculated that the density could stem from interactions involving the CTD.

To test this possibility, we turned to AlphaFold (AF), a neuronal network that can predict the three-dimensional structures of protein complexes with an accuracy that rivals experimentally determined structures ^30,31^. Capitalizing on the ability to AF to dock short, disordered peptide segments with high confidence ^32^, we divided the IDR of NPM1 into peptides of 20 amino acids with a 10-residue overlap and predicted possible complexes with the CTD (Figure 3e, Figure S5). We then calculated the binding energies for the top-scoring complex for each peptide. We found that peptides covering the acidic tracts A2 or A3 exhibited weakly favorable interactions due to charge contacts with the basic residues in the CTD. Placement of the top-scoring complex between the CTD and the A2 tract into the EM map showed a good agreement between the unassigned density, the helical acidic region of the IDR, and the approximate location of the CTD (Figure S4d). Considering the flexibility of the IDR, the CTD would be unlikely to occupy a more specific orientation, giving rise to a more diffuse density than the pentameric NTD.

To confirm the interaction between the disordered region and CTD, we recorded mass spectra of NPM1_188_ and NPM1_240-294_. We observed unresolvable peaks and a shift to the higher *m/z* region for NPM1_188_, indicating binding of the CTD with a mixed stoichiometry, as well as potentially inducing oligomerization (Figure S6a). Next, we designed a short NPM1 variant, NPM1_βIDR_, in which NTD and CTD are linked without the IDR (Figure 3f). Native MS analysis of NPM1_βIDR_ revealed almost exclusively pentameric protein and a small decamer population, irrespective of collision voltage (Figure 3f). This result demonstrates that removing the IDR prevents higher oligomerization of NPM1 pentamers. Taken together, fluorescence spectroscopy, AF, cryo-EM, and MS suggest that the CTD engages in weak, “fuzzy” interactions with the disordered region of neighboring pentamers. The IDR and CTD can be viewed as a “grappling hook” that links NPM1 into large oligomers.

To investigate how the interactions between CTD and IDR affect nucleotide binding to NPM1, we recorded mass spectra of FL NPM1 in the presence of equimolar amounts of tRNA, which binds to NPM1 and is more homogeneous than rRNA and thus easier to detect by MS. We found that tRNA completely abolishes the NPM1 signal in MS even at high collision voltages (Figure S6b), consistent with tRNA cross-linking the

NPM1 pentamers into oligomers that are too stable for gas-phase dissociation. We also considered AML mutations that disrupt the third helix of the CTD and asked whether they may impact self-assembly. We purified an NPM1 variant with a mutation at the C-terminus (NPM1_AML_) in which the last seven residues (WQWRKSL) are exchanged for an eleven-residue sequence lacking the nucleolar localization signal (CLAVEEVSLRK) and recorded mass spectra under native conditions. Strikingly, we did not observe a pronounced difference between FL NPM1 and NPM1_AML_ (Figure S6c). We conclude that the AML variants do not exhibit altered self-assembly properties, and that its oncogenic potential can therefore be attributed to loss of the nucleolar localization signal in helix 3.

### Nucleolar components modulate NPM1 self-assembly and chaperone activity

Having established that CTD interactions mediate NPM1 self-assembly, we then asked whether the same interactions affect its ability to delay Aβ_42_ fibrillation. It is well-established that NPM1 can undergo LLPS through heterotypic interactions with RNA or basic peptides located in the nucleolus. Peptides such as residues 21-37 of ribosomal protein L5 (rpL5) or residues 299-326 of SURF6, which both contain multiple arginine-rich motifs, bind to the A1 tract of NPM1 and induce LLPS ^13^. The same type of interaction has been found to mediate sequestration of p14^ARF^ by nucleolar NPM1 ^33,34^. We speculated that such interactions would compete with higher oligomerization via the CTD. To test this possibility, we selected rpL5_21-37_, a short peptide with two basic motifs and whose association with NPM1 has been investigated in detail (Figure 4a) ^13,35^. The AF model shows binding of the second basic motif of rpL5_21-37_ to the acidic A1 groove on the NTD, while the first basic motif is extended away from the protein, making it accessible for charge interactions with another NTD (Figure 4a). This model, in agreement with previous reports ^13^, thus suggests that rpL5_21-37_ cross-links acidic regions in NPM1. We monitored the effect of rpL5_21-37_ on NPM1 self-assembly by incubating FL NPM1 for 60 min at room temperature in the absence or the presence of increasing concentrations of the peptide and following the accumulation of sedimented macroscopic assemblies by light microscopy (Figure S7a). After incubation of NPM1 alone, we detected a small number of spherical aggregates, which may be related LLPS of NPM1 in the presence of trace amounts of nucleotides not detectable by MS. Addition of rpL5_21-37_ at a ratio of 1:10 caused a notable increase in assembly size and fusion into larger, amorphous structures. After 60 min incubation at a 10-fold excess of rpL5_21-37_, exclusively amorphous assemblies were observed. Since rpL5_21-37_ can induce the assembly of both FL NPM1 as well as the isolated NTD ^13^, we recorded mass spectra of NPM1_120_ in the presence of rpL5_21-37_ (Figure 4b). We observed significant peak broadening, indicating binding of the 2.2 kDa peptide to NPM1_120_ oligomers, as well as a shift from predominantly decamers to oligomers composed of ≥ 25 subunits. Next, we subjected complexes between FL NPM1 and rpL5_21-37_ to MS analysis. Notably, we did not detect pentamers or their fragments upon collisional activation, but only trace amounts of monomers (Figure 4c). Together, the insights from MS suggest that rpL5_21-37_ binding increases the size and stability of NPM1 oligomers by cross-linking acidic tracts. Importantly, these oligomers have a considerably closer association between NTD pentamers than those formed by interactions with the CTD, due to the short length of the rpL5_21-37_ peptide.

**Figure 4.**
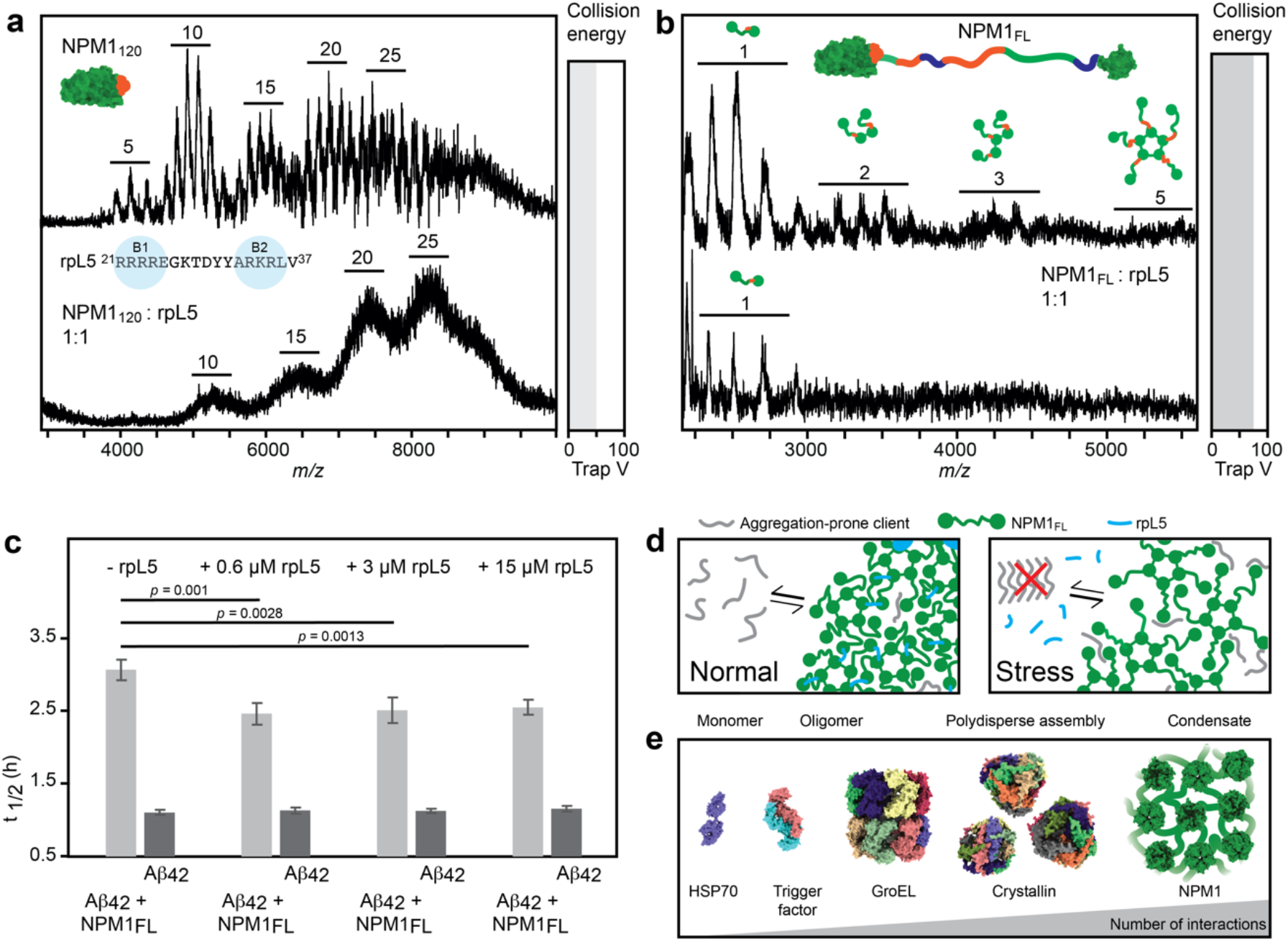
Modulating NPM1 self-assembly impacts its chaperone activity. (a) Residues 21-37 of rpL5 contain two basic motifs (B1 and B2). The AF model of rpL5_21-37_ (shown as turquoise ribbon, K/R residues as sticks) bound to the NTD pentamer (shown as electrostatic surface rendering) suggests binding of basic motif 2 to the acidic groove in the A1 tract of NPM1, while basic motif 1 extends away from the protein and is available for additional interactions. (b) Native mass spectra of NPM1_120_ show peak broadening rpL5_21-37_ and a shift to higher oligomeric states in the presence of rpL5_21-37_. (c) FL NPM1 assemblies formed with rpL5_21-37_ cannot be dissociated by collisional activation, as judged by the low abundance of monomers and the absence of dimers, trimers, or pentamers in the mass spectra. (d) Comparing the t _1/2_ for fibril formation as judged by ThT fluorescence shows that rpL5_21-37_ alone does not affect Aβ_42_ aggregation, whereas rpL5_21-37_ reduces the ability of NPM1 to delay Aβ_42_ fibrillation. Error bars indicate the standard deviation of n=4 repeats. Significance was calculated using Student’s *t*-test for paired samples with equal variance. (e) Proposed connection between chaperone activity and self-assembly of NPM1. Under normal conditions, NPM1 (green) and basic proteins (blue) such as rpL5 form a tight nucleolar network. Under stress, rpL5 is released from the nucleolus, loosening up the NPM1 network, which enables NPM1 to sequester and chaperone amyloidogenic client proteins (grey). (f) Oligomerization and chaperones. Chaperone stoichiometries range from monomers (HSP70) to polydisperse oligomers (α-crystallin B). Chaperone activity of NPM1, on the other hand, involves the formation of large assemblies without defined stoichiometry.

The fact that the rpL5_21-37_ peptide modulates higher oligomerization of NPM1 enabled us to test the impact of self-assembly on chaperone activity. We, therefore, performed aggregation assays with Aβ_42_ and FL NPM1 in the presence of increasing amounts of rpL5_21-37_ (Figure 4d). Although rpL5_21-37_ alone had no effect on Aβ_42_ fibrillation, it significantly reduced the ability of NPM1 to delay the formation of ThT-positive aggregates (Figure 4d, Figure S7b). Interestingly, the effect was observed already at a five-fold excess of NPM1, corresponding to one rpL5_21-37_ per pentamer. These data suggest that inducing a tighter and more complete association of NPM1 subunits through basic nucleolar peptides reduces its chaperone activity. To investigate whether the observations for rpL5_21-37_ represent a general mechanism, we also tested the SURF6_299-326_ peptide, which encompasses basic tract 4 of SURF6 and induces LLPS of NPM1 ^13,36^. AF models indicate a binding mode similar to that of rpL5_21-37_ (Figure S7c), in good agreement with their comparable effects on NPM1 assembly ^13,36^. Aggregation assays showed that SURF6_299-326_ alone has no impact on Aβ_42_ fibrillation but reduces the efficiency of FL NPM1 chaperoning to a similar extent as rpL5_21-37_ (Figure S7d). We conclude that proteins or peptides with multiple basic motifs cross-link NPM1 oligomers, which reduces their ability to delay fibril formation *in vitro*.

## Discussion

The nucleolar scaffold protein NPM1 controls p53-dependent tumor suppression by sequestering aggregation-prone transcription factors, including p14^ARF^ and c-MYC ^7^ and is implicated in the formation of amyloid bodies under stress conditions ^18^. In this study, we have investigated the basis of NPM1 self-assembly and how it relates to its ability to prevent amyloid formation *in vitro*. The CTD of NPM1 can delay fibrillation of the model amyloid Aβ_42_, yet this chaperone activity is modulated by NPM1 self-assembly. Using a combination of native MS, machine learning, and cryo-EM, we show that direct interactions between CTD and the acidic IDR in NPM1 induce the formation of an NPM1 network and thus reduce its ability to delay Aβ_42_ aggregation. Our findings imply that NPM1 pentamers are linked by their CTDs and IDRs, although additional interactions of the CTDs with the acidic NTDs cannot be excluded, as AF models of 29 homologs from the Pfam with identical domain architecture as NPM1 with sequence identities below 50% show clustering of the CTDs around the acidic side of the NTDs (Figure S8). The length of the IDR (approximately 120 residues) leaves considerable space between the NPM1 pentamers, creating flexible compartments lined with acidic and some basic residues. Previous studies have suggested diffuse interactions between the acidic and basic tracts in oligomers ^13^, which could regulate the degree of compaction of the NPM1 network. The addition of basic peptides such as rpL5 or SURF6 cross-links the acidic NTDs, and potentially the other acidic tracts, to constrict the network, in line with the induction of LLPS of NPM1 by polycations ^9,13^ (Figure 4e).

Our findings raise the question of how exactly NPM1 oligomers interact with unfolded or aggregation-prone clients. The observation that NPM1 does not seem to specifically inhibit primary or secondary nucleation or fibril elongation, but rather affects all these events, indicates that NPM1 works by partially sequestering Aβ_42_. We speculate that a fraction of the Aβ_42_ species are trapped in the NPM1 network and released on an equilibrium basis. Importantly, Aβ_42_ has not been identified as a client of NPM1 *in vivo* and does not include highly charged motifs that would promote recruitment into the NPM1 network via interactions with basic or acidic tracts. The fact that the chaperone activity of the CTD is higher than that of FL NPM1 suggests that Aβ_42_ can interact with free CTDs, and that this interaction is reduced when the CTDs instead bind to IDRs of neighboring NPM1 pentamers. The artificial amyloid β17, which shares some characteristics with Aβ_42_, co-localizes with NPM1 in cells ^17^, underscoring the possibility that NPM1 may bind a wider range of aggregation-prone sequences. Importantly, the isolated CTD does not prevent aggregation of denatured globular proteins, which instead requires FL NPM1 ^11,15^. These differences indicate different mechanisms for chaperoning unfolded proteins and amyloidogenic peptides, a distinction that has been reported for other chaperone systems ^37–39^. Unfolded proteins expose hydrophobic as well as charged patches, which may drive their recruitment into NPM1 oligomers. The fact that the nucleolus does not actively promote refolding but holds unfolded proteins before transferring them to the HSP70 network supports this concept ^17^.

We hypothesize, based on our findings, that changes in nucleolar composition could balance the ability of the NPM1 scaffold to chaperone unfolded proteins or amyloids. Basic proteins including rpL5 and p14^ARF^ mediate tight connections between the NPM1 pentamers but are released under stress conditions. The resulting loosening of the NPM1 network serves to activate the chaperone function of NPM1, in line with its role as safeguard against stress-induced protein aggregation (Figure 4e). In this context, NPM1 may be viewed as an extreme case of a polydisperse chaperone (Figure 4f). Interestingly, a protein with similarities to NPM1 has recently been identified as a non-canonical chaperone: the death-domain associated protein (DAXX) contains a folded N-terminal domain as well as a disordered region with poly-glutamate and poly-aspartate stretches and undergoes LLPS through interactions with p62 ^40^. DAXX can prevent Aβ_42_ fibrillation as well as the aggregation of heat-denatured proteins and associates with p53 to ensure correct folding ^41^. It furthermore controls nucleolar integrity^42^, raising the possibility that poly-D/E proteins constitute a class of chaperones that incorporate into nucleolar condensates.

Although our study clearly shows that direct interactions involving the CTD modulate the chaperone activity of NPM1, it has some limitations. Specifically, the experimental approach used here (cryo-EM, AF, and native MS) is best-suited for detecting complexes between folded domains. For example, the density map outside of the NTD is too diffuse to allow any unambiguous reconstructions, and we could not observe by EM binding of the CTD to IDR further away from the NTD. Similarly, AF does not reliably predict preferred conformations of disordered regions. As a result, we could not obtain atomistic insights into complexes between the CTD and FL NPM1. More broadly, self-assembly of NPM1 is affected by multiple types of “fuzzy” interactions, including with polyanions, polycations, and molecular crowders ^9^, but we cannot independently assess their contributions to NPM1 chaperone activity within the scope of this study. Our results indicate that RNA and basic peptides increase the stability of NPM1 oligomers, but do not reveal whether this effect is related to phase separation of NPM1. For example, we observe a shift to larger oligomers of NPM_120_ in the presence of rpL5_12-37_. However, the isolated NTD does not undergo phase separation with rpL5_12-37_ unless the disordered A2 tract is included ^13^, which hints at a need for flexible connections between the protomers for LLPS to occur. Similarly, RNA molecules could increase the distance between NTD pentamers and thus modulate the fluidity of the assembly. Further studies are therefore needed to assess the specific role(s) of LLPS in nucleolar chaperoning.

## Experimental

### Protein sequences

Hexahistidine (H10) affinity tag with tobacco etch virus (TEV) cleavage (/) site MGHHHHHHHHHHENLYFQ/S

NPM1_FL_

UniProtKB - P06748 (NPM_HUMAN) Acidic tracts 1, 2, and 3 are shown in bold. Positions 120, 188, and 240 for NPM1_120_, NPM1_188_, and NPM1_240_ are indicated.

^1^MEDSMDMDMSPLRPQNYLFGCELKADKDYHFKV**DNDENE**HQLSLRTVSLGAGAKDELHI

VEAEAMNYEGSPIKVTLATLKMSVQPTVSLGGFEITPPVVLRLKCGSGPVHISGQHLVAV**E**^120^

**EDAESEDEEEED**VKLLSISGKRSAPGGGSKVPQKKVKLAA**DEDDDDDDEEDDDEDDDDD**

**DFDDEEAEE**^188^KAPVKKSIRDTPAKNAQKSNQNGKDSKPSSTPRSKGQESFKKQEKTPKTP

KG^240^PSSVEDIKAKMQASIEKGGSLPKVEAKFINYVKNCFRMTDQEAIQDLWQWRKSL^294^

NPM1_AML_

Mutated residues are underlined.

^1^MEDSMDMDMSPLRPQNYLFGCELKADKDYHFKVDNDENEHQLSLRTVSLGAGAKDELHIVEAEAMNYEGSPIKVTLATLKMSVQPTVSLGGFEITPPVVLRLKCGSGPVHISGQHLVAVEEDAESEDEEEEDVKLLSISGKRSAPGGGSKVPQKKVKLAADEDDDDDDEEDDDEDDDDDDFDDEEAEEKAPVKKSIRDTPAKNAQKSNQNGKDSKPSSTPRSKGQESFKKQEKTPKTPKGPSSVEDIKAKMQASIEKGGSLPKVEAKFINYVKNCFRMTDQEAIQDLCLAVEEVSLRK^298^

NPM1_βIDR_

MEDSMDMDMSPLRPQNYLFGCELKADKDYHFKVDNDENEHQLSLRTVSLGAGAKDELHIV

EAEAMNYEGSPIKVTLATLKMSVQPTVSLGGFEITPPVVLRLKCGSGPVHISGQHLVAVE_P

SSVEDIKAKMQASIEKGGSLPKVEAKFINYVKNCFRMTDQEAIQDLWQWRKSL

### Protein expression and purification

The genes for MGH10-TEV-NPM1 and its corresponding mutants in the pET26b(+) expression vector were transformed into chemically competent *E. coli* BL21 (DE3) cells. Overnight cultures were inoculated 1:100 to Luria-Bertani medium containing 50 µg/mL kanamycin. The cultures were grown at +37°C to an OD600 of 0.6-0.9 before induction with 0.5 mM isopropyl-β-D-thiogalactopyranoside (IPTG) and overnight expression at +25°C. For each construct, 500-1000 mL of culture was harvested by centrifugation at 6000 × g for 20 min at RT. The pellet was resuspended in 20 mM Tris, pH 8.0 buffer (1 mL for 10 mL of culture) and stored at -20°C overnight. After thawing, the cells were lysed with the addition of 1 mg/mL lysozyme, 10 μg/ml DNAse I and 2 mM MgCl_2_ and incubated for 1 hour on ice followed by sonication using a probe sonicator (Qsonica, CT) on ice for 5 min, 0.5 s on/1 s off at 30% power. The samples were centrifugated for 15 min at 15 000 × g, 4°C, the supernatant was decanted, and the pellet was resuspended in 20 mM Tris, 500 mM NaCl pH 8 buffer (1 mL per 10 mL culture) sonicated and centrifuged as before. Two further resuspensions with the same parameters were conducted. Samples taken from steps during expression, lysis and purification were analyzed using SDS-PAGE 4-20% Mini-PROTEAN® TGX™ polyacrylamide gels (Bio-rad Laboratories Inc., USA) stained with Coomassie G-250 Brilliant Blue dye. The protein-containing supernatants were supplemented with 20 mM imidazole and 500 mM NaCl if needed, loaded onto 1 mL HiTrap IMAC HP (Cytiva, Sweden) columns charged with Co^2+^ and eluted with a linear gradient of buffer containing 500 mM imidazole, 500 mM NaCl, 20 mM

Tris, pH 8. Fractions were analysed using SDS-PAGE gels, protein-containing fractions were pooled, concentrated using appropriate MWCO Amicon ® Ultra 15 centrifugation tubes (Merck, USA) and dialysed at +4°C five times against 20 mM Tris, 500 mM NaCl, pH 8 using Slide-A-Lyzer™ MINI Dialysis Devices with 10K MWCO (Thermo Scientific, USA). Protein concentrations were determined using a Pierce™ BCA assay (Thermo Scientific, USA).

### Aβ_42_ aggregation assays

Aβ_42_ was produced as described previously ^43^. Aggregation kinetics were monitored in solution by measuring total ThT fluorescence using a POLARstar Omega microplate reader (BMG Labtech, Germany) with an excitation filter at 440 nm and an emission filter at 480 nm. All measurements were conducted at +37°C for 24 hours, without agitation. Samples were prepared with 10 μM ThT in 170 mM NaCl, 17 mM Tris, 3 mM NaPi, 0.03 mM EDTA pH 8 in black half-area 384-well polystyrene microplates with a transparent (Corning, USA). The total reactant volume of each replicate was 20 μl. For kinetic experiments with constant Aβ42 concentrations, a final concentration of 3 μM Aβ42 was prepared in the absence and presence of varying molar ratios of NPM1 variants. To determine the effects of different NPM1 concentrations on the aggregation half time of Aβ42 fibril formation, fluorescence data was normalized and fitted to an empirical logistic5 function with an equation (1):

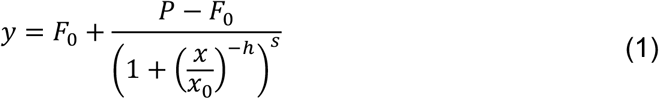

where F_0_ is the baseline value, P the plateau value, x the time point value, x_0_ the time factor, h the Hill slope steepness factor, s the control factor. The aggregation half-time τ_0_ values were derived from the following equation (2):

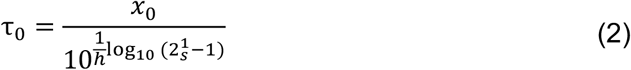

Where x_0_ is the time factor, h the Hill slope steepness factor, s the control factor. Data is represented as the average±standard deviation from 4-5 experiments.

### Native mass spectrometry

All purified NPM1 variants were buffer-exchanged into 1 M ammonium acetate, pH 8.0, using Bio-Spin P-6 columns (BioRad, CA). Mass spectra were acquired on a Waters Synapt G1 travelling wave ion mobility mass spectrometer modified for high-mass analysis (MS Vision, NL) equipped with an offline nanospray source. The capillary voltage was 1.5 kV, the source pressure was 8 mbar, and the source temperature was 80°C. Mass spectra were visualized using MassLynx 4.1 (Waters, UK).

### AF structure predictions

All AF predictions for NPM1 from *Homo sapiens* were generated using Colab Fold (https://colab.research.google.com/github/sokrypton/ColabFold/blob/main/AlphaFold2.ipynb) Version 1.4 with default settings (5 models, no AMBER step). Inclusion of an AMBER step did not yield noticeably different structures, as did inclusion of the polyhistidine-tag in the NPM1 sequence. Structures were visualized with ChimeraX V1.3 ^44^.

### Negative stain electron microscopy

For negative stain EM, a final concentration of 3 μM Aβ_42_, 3 μM NPM1_240-294_, or 3 μM Aβ_42_ + 3 μM NPM1_240-294_ were prepared in 160 mM NaCl, 16 mM Tris, 4 mM NaPi, 0.04 mM EDTA pH 8 and incubated overnight at 37°C. The samples were then centrifuged at 17000 x *g* for 30 min and 80% of the supernatant was removed. The remainder was resuspended and 5 µL was loaded to 200 mesh copper grids with Formvar/Carbon support film that had been glow-discharged at 25 mA for 2 min. After one minute of incubation, the liquid was removed and the negative staining was carried out by applying a 5 µl of 1 % (w/v) uranyl acetate (in H_2_O) to the grid for 20-30 seconds. Then, the liquid was removed and the procedure was repeated 6 times ^45^. The grids were imaged in Talos 120 C G2 (Thermo Scientific) equipped with a CETA-D detector.

### Specimen preparation for cryo-EM

Four microliters of 0.75 mg/ml of purified NPM1 were applied to Cryomatrix 2/1 grids (glow discharged for 2 min at 25 mA) in a Vitrobot MK IV (Thermo Fisher Scientific) at 4°C and 100% humidity. Sample excess was removed by blotting for 4 s using a blot force of 1 followed by vitrification in liquid ethane.

The data were collected on a Krios G3i electron microscope (Thermo Fisher Scientific) at an operating voltage of 300 kV with Gatan BioQuantum K3 image filter/detector (operated with a 10 eV slit) at the Karolinska Institutet’s 3D-EM facility, Stockholm, Sweden. The data were collected using EPU (Thermo Fisher Scientific). An EFTEM SA magnification of 165,000x was used, resulting in a pixel size of 0.505 Å, with a total dose of 54 e^-^/Å^2^ divided across 60 frames over 2 s (fluency of 6.9 e-/px/sec). Target defocus values were set between -0.2 to -2 µm and using a stage tilt of 0° or -20°. The data processing strategy is schematized in Fig S3a. Motion correction and CTF estimation was performed in Warp ^46^.The micrographs were then imported in Scipion3 ^47^ and denoised using Janni for picking with crYOLO ^48^ using the pre-trained model for de-noised micrographs. The particle picks were pruned using XMIPP Deep Micrograph Cleaner ^49,50^. The particles were then extracted in WARP (64 px box, 2-fold binning to 2.02 Å/px). 2D classification of the particle set was performed in CryoSPARC v3.01 ^51^ and the good 2D classes were manually selected for *ab-initio* reconstruction in CryoSPARC of 1 class. The particles were then refined in RELION 4.0 beta ^52^.The refined particles were re-extracted in WARP (192 px box, 1.01 Å/px) and further refined in RELION 4.0 beta using a mask (covering N-terminal and C-terminal domains) and subjected to 3D classification without alignment (6 classes, T 20) using the same mask. The class with very weak density for C-terminal domains (Class 1) and the class with the strongest density for C-terminal domains (Class 6) were then selected and refined individually in CryoSPARC v3 non-uniform refinement. In order to improve the resolution on the N-terminal region and be able to fit an atomic model for this region, Class 6 was further subjected to 3D classification in RELION 4.0 beta. The two classes which reconstructed to the highest resolution were pooled and re-extracted in WARP (382 px box, 0.505 Å/px). These particles (106,905 particles) were finally refined and post-processed in RELION 4.0 with a mask on the N-terminal region. The resulting map was used for the refinement of an atomic model for the N-terminal region. The PDB 5EHD, which was used as a starting model, was docked in the map and refined in Coot 0.9.8.1 ^53^ and REFMAC5 ^54^. Refinement data for Table S1 was generated with MolProbity ^55^ in Phenix 1.20.1 ^56^

### Binding energy calculations

The peptide: protein binding energy was computed using MM/GBSA (Molecular Mechanics / Generalized Born Surface Area) method ^57^ as obtained from the following equations:

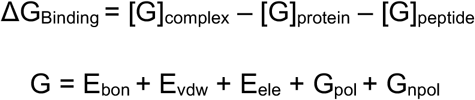

E_bon_ + E_vdw_ + E_ele_ + G_pol_ + G_npol_E_bon_ is composed of three bonded (bond, angle and dihedral) energy terms. E_vdw_ and E_elec_ are the van der waals and electrostatic non-bonded interaction components respectively. These energies are calculated using Molecular Mechanics (MM) force field expressions of AMBER. G_pol_ is the polar solvation energy which is obtained by solving the Generalized Born (GB) solvation model. G_npol_ is the non-polar solvation energy estimated through an empirical linear relationship (γ*SASA), where “γ” is the surface-tension and “SASA” is the Solvent Accessible Surface Area. The complex state structures were used to generate the corresponding free states of the protein and peptide. The continuum solvent environment was represented with an implicit GB model (IGB=2). The internal dielectric constant for the protein/peptide and the solvent was set to 1 and 80 respectively, γ= 0.0072 kcal/mol/Å^2^ and the salt-concentration was set to 0.15 mM. The MMPBSA.py script ^58^ available through the AMBER18 suite of programs was used to carry out the calculations.

### Light microscopy

Unlabeled NPM1 at 10 μM in 150 mM NaCl, 2 mM DTT, in 10 mM Tris pH 7.5 with or without basic peptides at concentrations of 10 or 100 μM were added to Corning 3651 polystyrene 96-well plates (Corning, USA). The mixtures were incubated at room temperature for 1 h and the liquid-liquid phase separation images in brightfield mode were recorded using a Zeiss Cell Observer microscopy instrument (Carl Zeiss AG, Germany) at a final magnification of 400x using LD Plan Neofluor 0.6 Corr Air objective. Images were recorded from initial focus point ±40 μm with a step of 1 μm.

### Tryptophan fluorescence spectroscopy

c-MYC *Pu24I* Quadruplex DNA (TGA GGG TGG IGA GGG TGG GGA AGG) was diluted in 10 mM Tris, 100 mM KCl pH 7.4 to a final concentration of 100 μM. The quadruplex DNA was formed by denaturation at +95°C for 15 min followed by overnight cooling to room temperature in a heating block. For tryptophan fluorescence spectroscopy, the samples were incubated for 2 h at +37°C on a Tecan Spark 20M multimode reader (Tecan Instruments, Switzerland) using Corning 3642 polystyrene 96-well plates (Corning, USA) and then excited at 280 nm (bandwidth 10 nm) with emission recorded at 350 nm (bandwidth 10 nm). The gain was set automatically, and data were analyzed with the Magellan software package (Tecan Instruments, Switzerland). NPM1_240_, NPM1_240-294_ and quadruplex DNA were tested at concentrations of 30 μM each. Results are expressed as average±standard deviation of three replicates.

### Competing interest statement

The authors declare no competing interests.

## Data availability

The density maps for the N-terminal region (”CryoEM structure of the human Nucleophosmin 1 core”) and for the two selected 3D classes have been deposited in the EMDB (EMD-15606). The refined atomic coordinates for the NPM1 core have been deposited in the PDB with (code 8AS5). All other data are available from the corresponding authors upon reasonable request.

## Author contributions

MS and ML designed the study with input from TMA, MAH, DPL, and JJ. MS and MK produced NPM1 variants and GC produced Aβ_42_. MS and AL performed aggregation assays. MS and CS performed light microscopy analysis. DL performed energy calculations. GVG and MH recorded and analyzed EM data. ML and AE generated structure predictions. ML wrote the paper with input from all authors.

## Acknowledgements

ML is supported by a KI faculty-funded Career Position, a KI-Cancer Blue Sky grant, a Cancerfonden Project grant (19 0480), and a VR Starting Grant (2019-01961). AE is supported by the Olle Engkvist Foundation (214-0344, to ML), CS is supported by a Novo Nordisk Foundation Postdoctoral Fellowship (NNF19OC0055700). DPL is supported by a Swedish Research Council grant for Internationally Recruited Scientists (2013-08807). AE is supported by Swedish Research Council for Natural Science, grant No. VR-2016-06301, Swedish E-science Research Centre, and the Knut and Alice Wallenberg Foundation. Computational resources: Swedish National Infrastructure for Computing, grants: SNIC 2021/5-297, SNIC 2021/6-197 and Berzelius-2021-29. The authors extend special thanks to Sergej Masich and access to the KI 3D-EM facility. ML gratefully acknowledges technical support from MS Vision, NL.

## Notes

### Competing Interest Statement

The authors have declared no competing interest.

