## Supplementary material for "A “grappling hook” interaction balances self-assembly and chaperone activity of Nucleophosmin 1": Table S1

**Table S1. Model and data statistics**

The data was generated with MolProbity in Phenix 1.20.1

| | |
|---|---|
| Model | |
| Composition (#) | |
| Chains | 5 |
| Atoms | 3673 (Hydrogens: 0) |
| Residues | Protein: 481 Nucleotide: 0 |
| Water | 0 |
| Ligands | 0 |
| Bonds (RMSD) | |
| Length (Å) (# > 4sigma) | 0.007 (0) |
| Angles (°) (# > 4sigma) | 1.193 (0) |
| MolProbity score | 1.84 |
| Clash score | 11.75 |
| Ramachandran plot (%) | |
| Outliers | 0.22 |
| Allowed | 1.56 |
| Favored | 98.22 |
| Rama-Z (Ramachandran plot Z-score, RMSD) | |
| whole (N = 449) | 0.39 (0.36) |
| helix (N = 0) | --- (---) |
| sheet (N = 198) | 1.48 (0.35) |
| loop (N = 251) | -0.82 (0.33) |
| Rotamer outliers (%) | 2.18 |
| Cbeta outliers (%) | 0 |
| Peptide plane (%) | |
| Cis proline/general | 37.5/0.0 |
| Twisted proline/general | 0.0/0.0 |
| CaBLAM outliers (%) | 2.64 |
| ADP (B-factors) | |
| Iso/Aniso (#) | 3673/0 |
| min/max/mean | |
| Protein | 42.43/302.83/130.98 |
| Nucleotide | --- |
| Ligand | --- |
| Water | --- |
| Occupancy | |
| Mean | 1 |
| occ = 1 (%) | 100 |
| 0 < occ < 1 (%) | 0 |
| occ > 1 (%) | 0 |
| Data | |

| | | |
|---|---|---|
| Box | | |
| Lengths (Å) | 71.71, 69.69, 60.09 | |
| Angles (°) | 90.00, 90.00, 90.00 | |
| Supplied Resolution (Å) | 2.5 | |
| Resolution Estimates (Å) | Masked | Unmasked |
| d FSC (half maps; 0.143) | 3.5 | 3.6 |
| d 99 (full/half1/half2) | 2.1/1.0/1.0 | 1.3/1.0/1.0 |
| d model | 3.5 | 3.6 |
| d FSC model (0/0.143/0.5) | 2.2/2.8/3.7 | 2.2/3.1/3.8 |
| Map min/max/mean | -0.13268E-02/0.42573E-02/0.63599E-05 | |
| Model vs. Data | | |
| CC (mask) | 0.66 | |
| CC (box) | 0.8 | |
| CC (peaks) | 0.56 | |
| CC (volume) | 0.64 | |
| Mean CC for ligands | --- | |

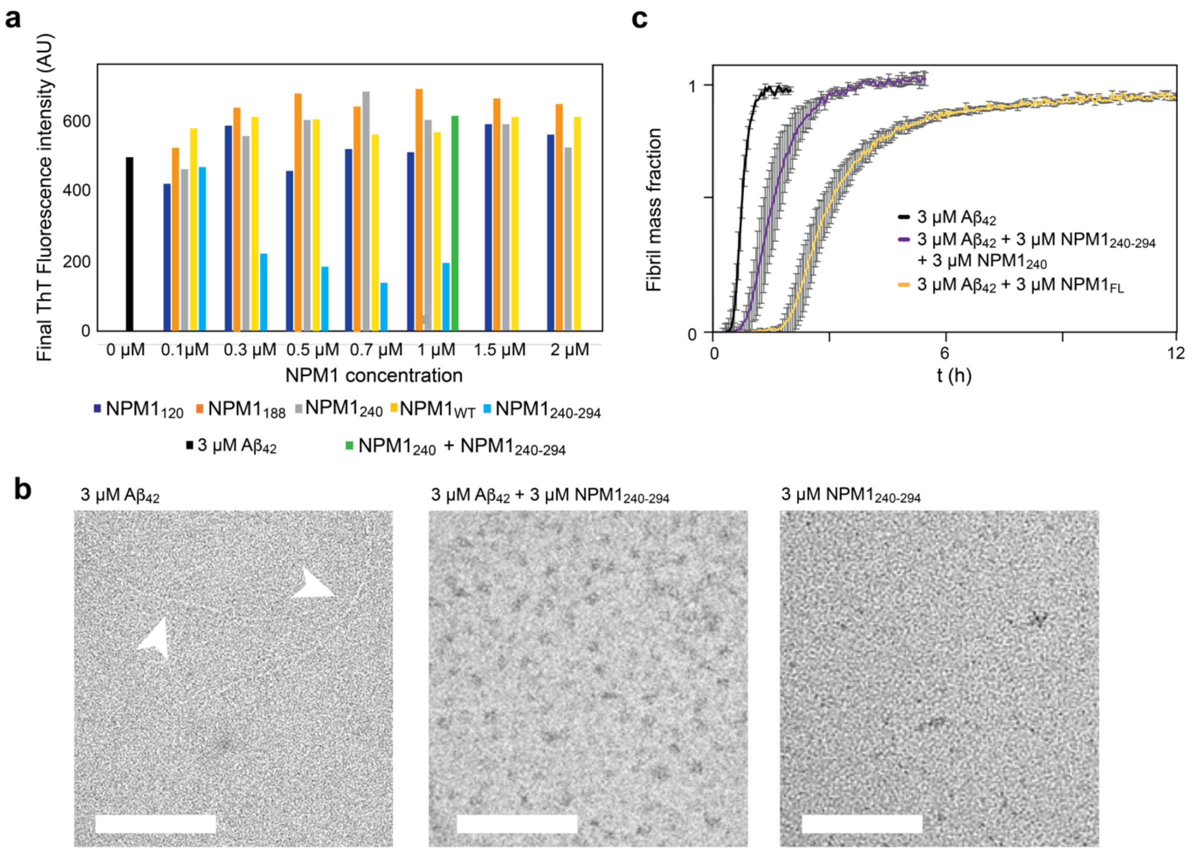


**Figure S1. Effect of $NPM_{240\text{-}294}$ on $A\beta_{42}$ aggregation.** (a) Final ThT fluorescence intensities of $A\beta_{42}$ for all NPM1 constructs shown in Figure 1 indicate similar amounts of fibrils formed, except for incubation with $NPM_{240\text{-}294}$. (b) Negative stain electron microscopy of the reaction endpoints (overnight) of $A\beta_{42}$ alone (left), $A\beta_{42}$ in the presence of $NPM_{240\text{-}294}$ (middle) and $NPM_{240\text{-}294}$ alone (right). Fibrils are observed only for $A\beta_{42}$ alone (white arrows). For $A\beta_{42}$ and $NPM_{240\text{-}294}$, some amorphous aggregates can be detected. Scale bars are 100 nm. (c) ThT fluorescence curves for aggregation of $A\beta_{42}$ in the presence of $NPM1_{240}$ and $NPM1_{240\text{-}294}$ show that the addition of the NTD reduces chaperone activity of the CTD, also compared to FL NPM1. (b)

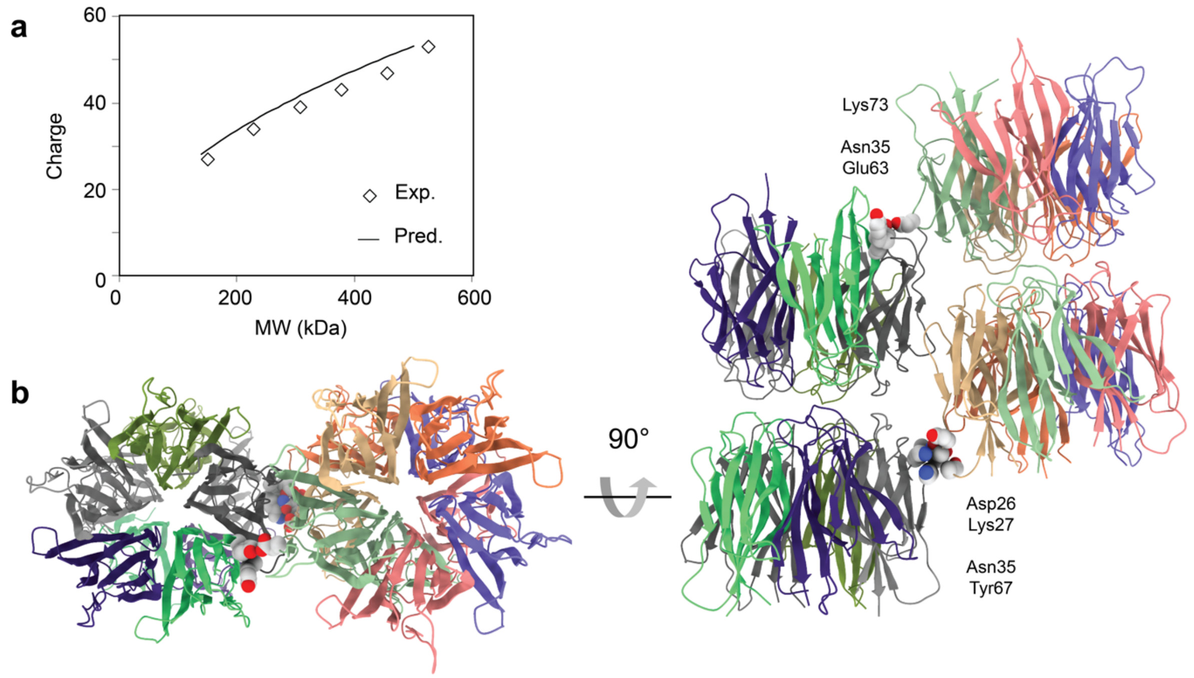


**Figure S2. Ordered oligomerization of $NPM_{120}$.** (a) Plotting the average charge of $NPM_{120}$ oligomers as a function of molecular weight reveals a good correlation with the charge expected for globular protein (solid line). (b) Crystal packing of NPM1 NTDs shows end-to-end pentamers making salt bridges with neighboring oligomers (PDB ID 5EHM). Similar contacts could mediate multimerization of $NPM_{120}$ observed by MS. Note stabilization of the acidic A1 tract (residues 34-39) through crystal contacts.

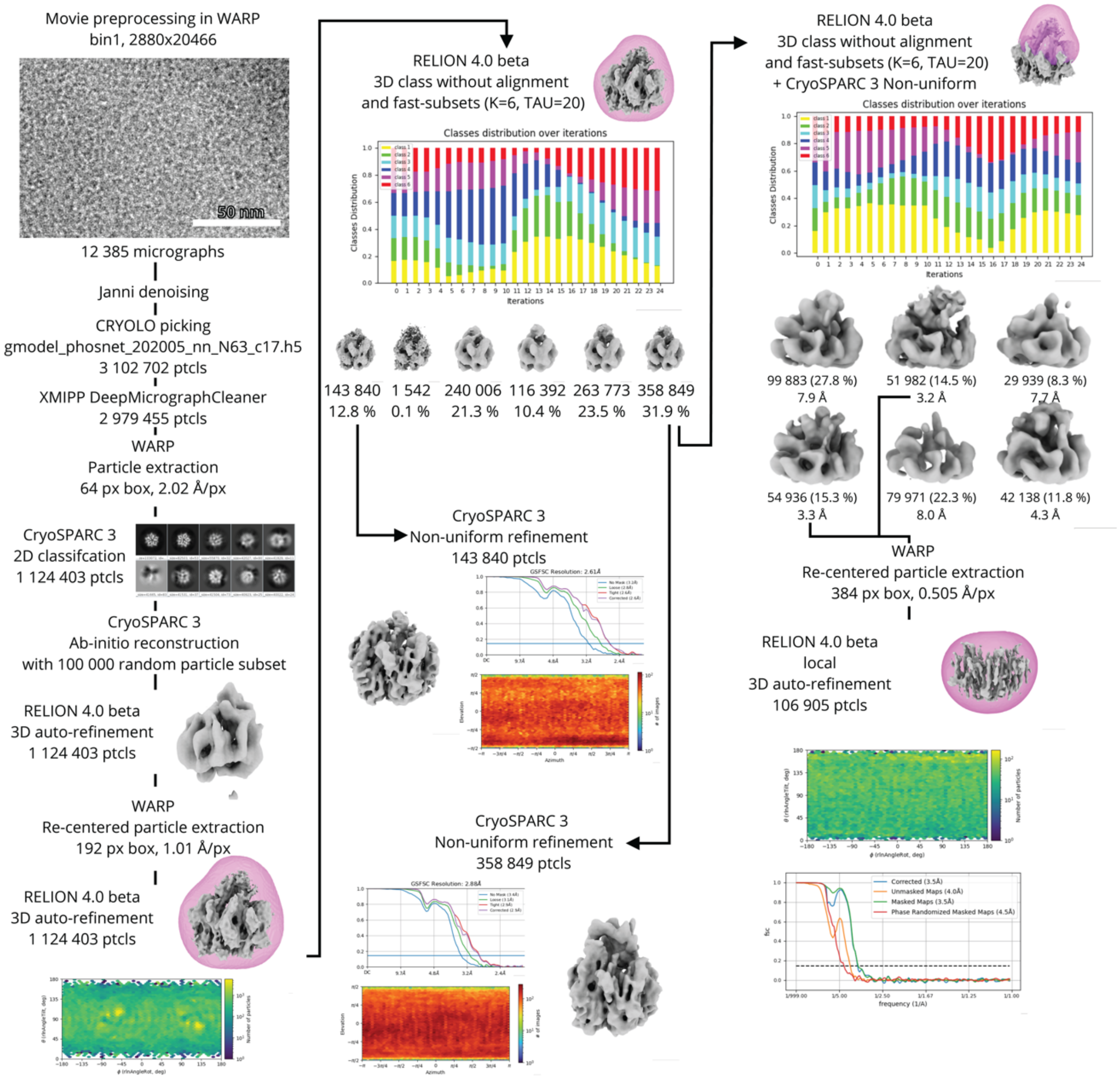


**Figure S3. Cryo-EM analysis strategy for FL NPM1.** The data analysis and density reconstruction strategy for NPM1.

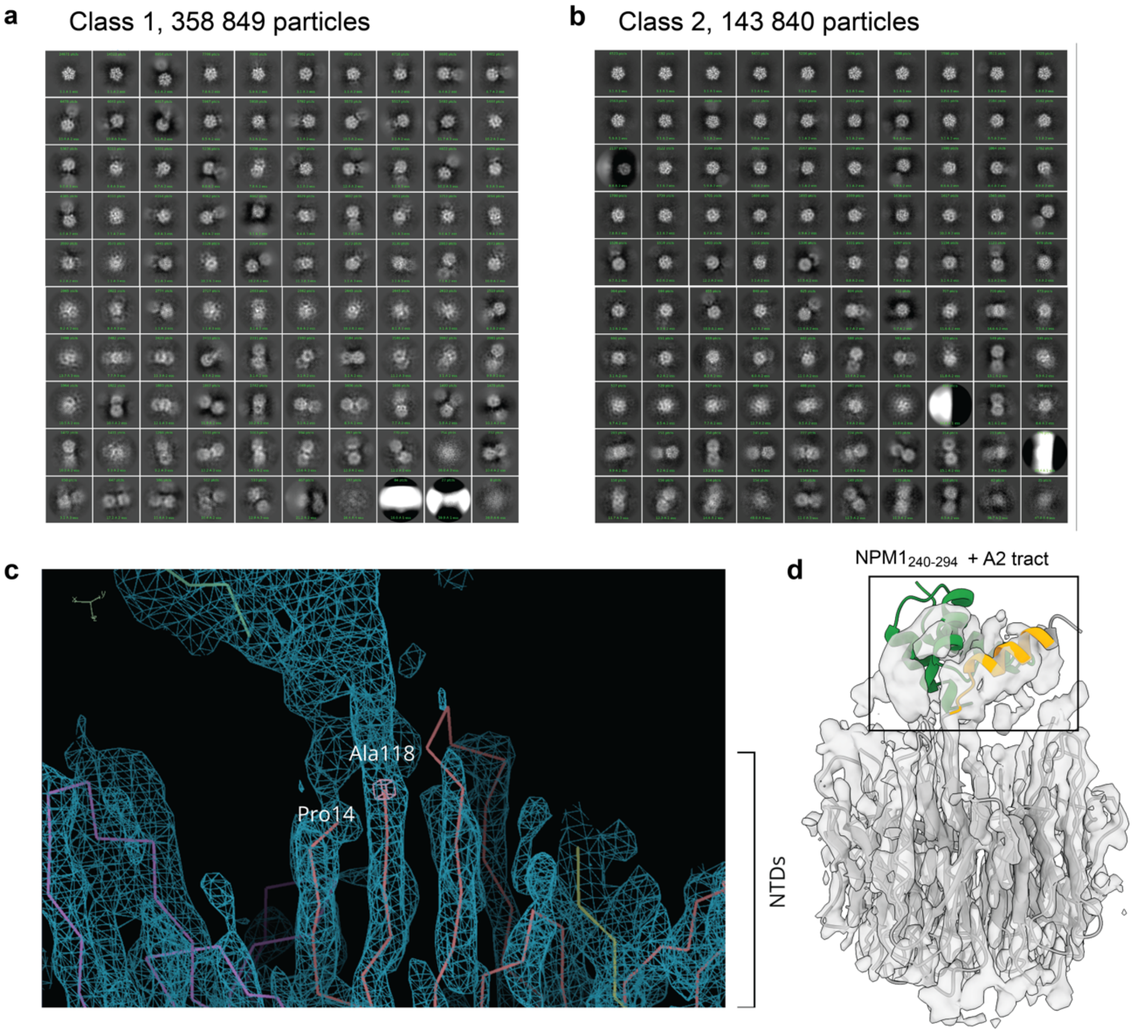


**Figure S4. Cryo-EM analysis of NPM1 particles.** (a) Particles in class 1 nearly always include one or two nearby particles that correspond to other NTD pentamers. (b) Particles in class 2 rarely have neighboring particles. (c) Fitting of the class 1 density map shows that the extra density above the NTD is connected to one protomer via its A2 tract, starting at residue 119. (d) Placement of the top-scoring CTD-A2 complex shows reasonable agreement between the location of the A2 tract and the CTD with the additional density in class 1 particles. The A2 tract, connected to the NTD, is shown in orange, the CTD in green. However, the A2-CTD complex can be fitted in multiple positions.

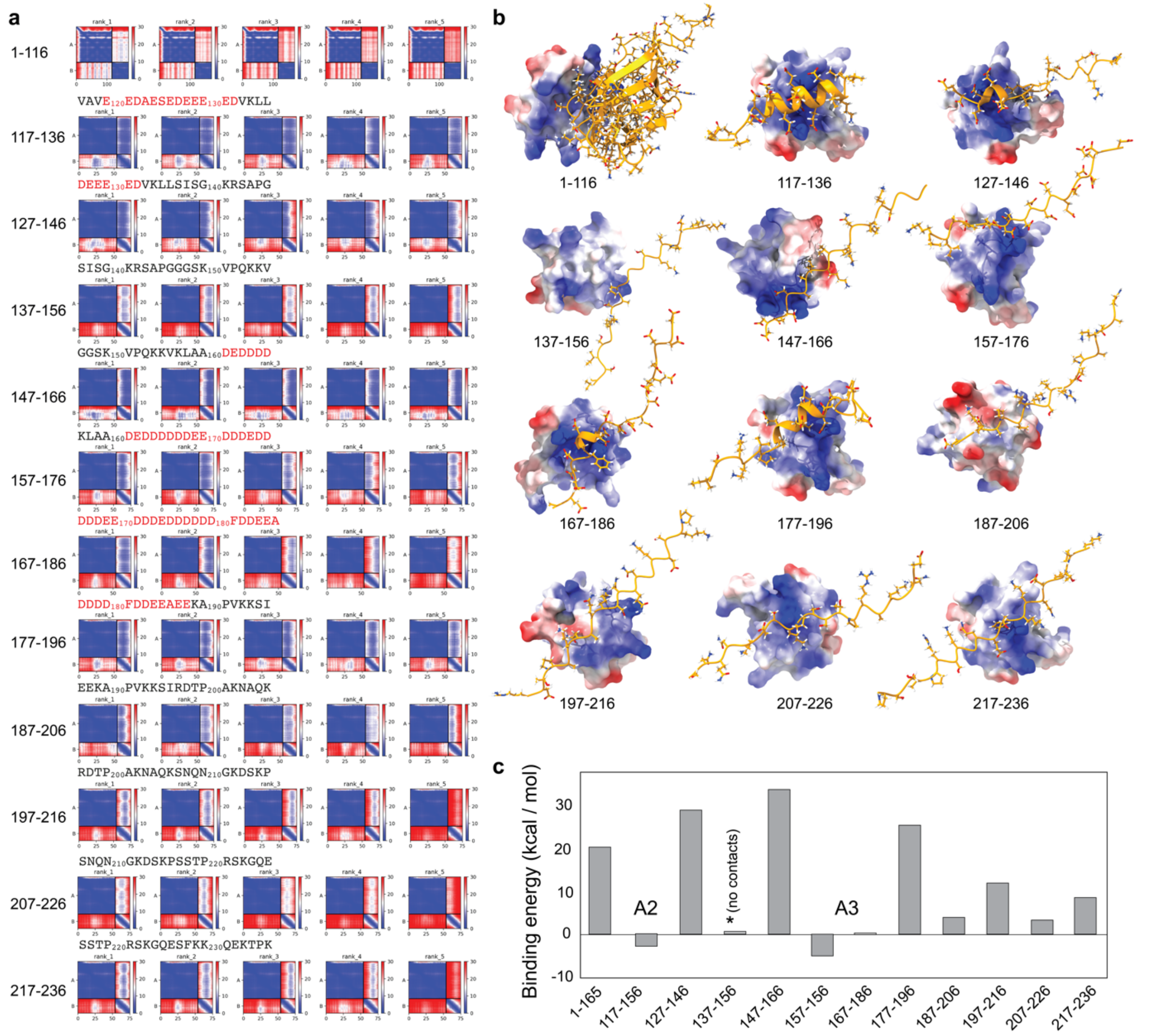


**Figure S5. AF docking of the NTD and overlapping peptides covering the IDR to the CTD.** (a) PAE plots show well-defined complexes between the CTD and peptides covering the A2 or A3 tracts. (b) The top-scoring models for each CTD-peptide complex are shown as electrostatic surfaces (CTD) and organe ribbons (peptides). Note that residues 136-154 make no contacts with the CTD. (c) Free energy calculations show weakly favorable interactions between the A2 and A3 tracts and the CTD.

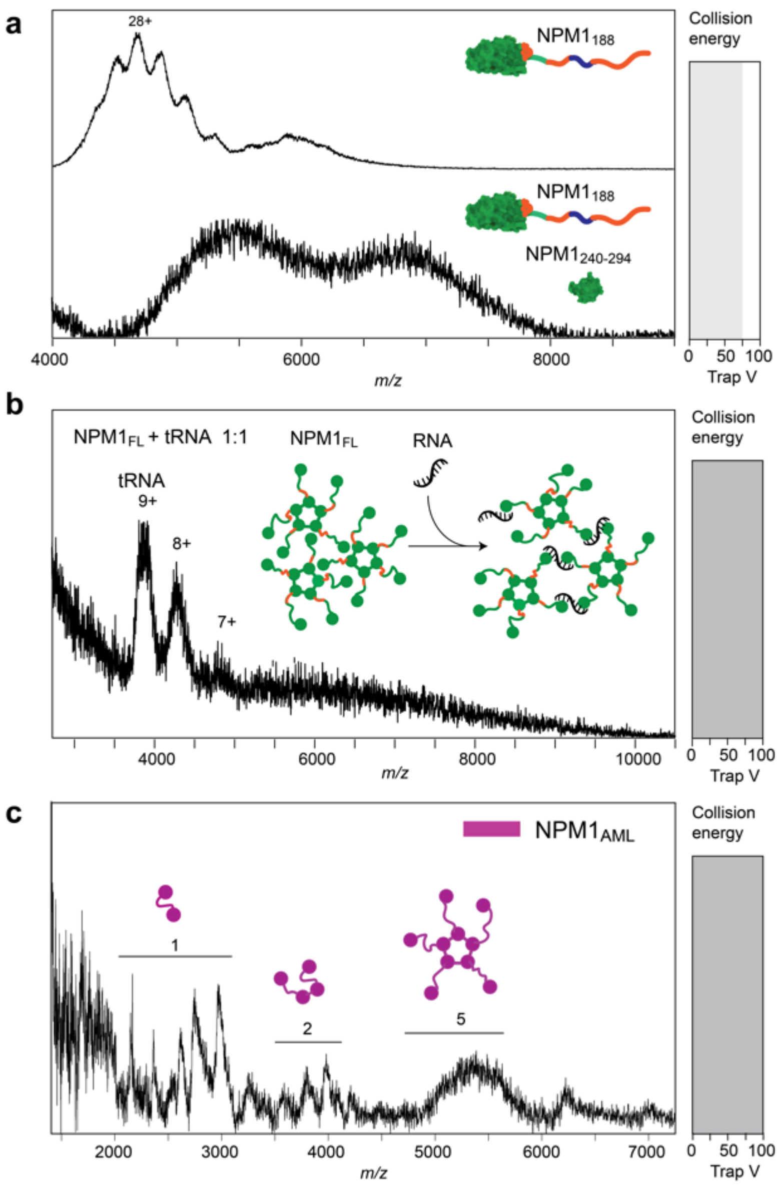


**Figure S6. Interactions of wt and mutant NPM1 studied by MS.** (a) Native MS of $NPM1_{188}$ in the absence (top) or the presence (bottom) of $NPM1_{240\text{-}294}$ shows a shift to the higher *m/z* region as well as significant peak broadening. (b) Addition of an equimolar amount of tRNA to NPM1 results in complete absence of NPM1 signal in mass spectra, consistent with cross-linking of NPM1 by tRNAs into stable oligomers. (c) Native MS and AF predictions of NPM1 lacking residues 284-294 in the CTD. Native mass spectra at a collision voltage of 100 V show Pentamers and smaller oligomers, comparable to FL wild-type NPM1. The AF models of the CTD of the AML variant (pink) show only minor changes in binding to the A1 and A2 tracts compared to the wild-type CTD (green).

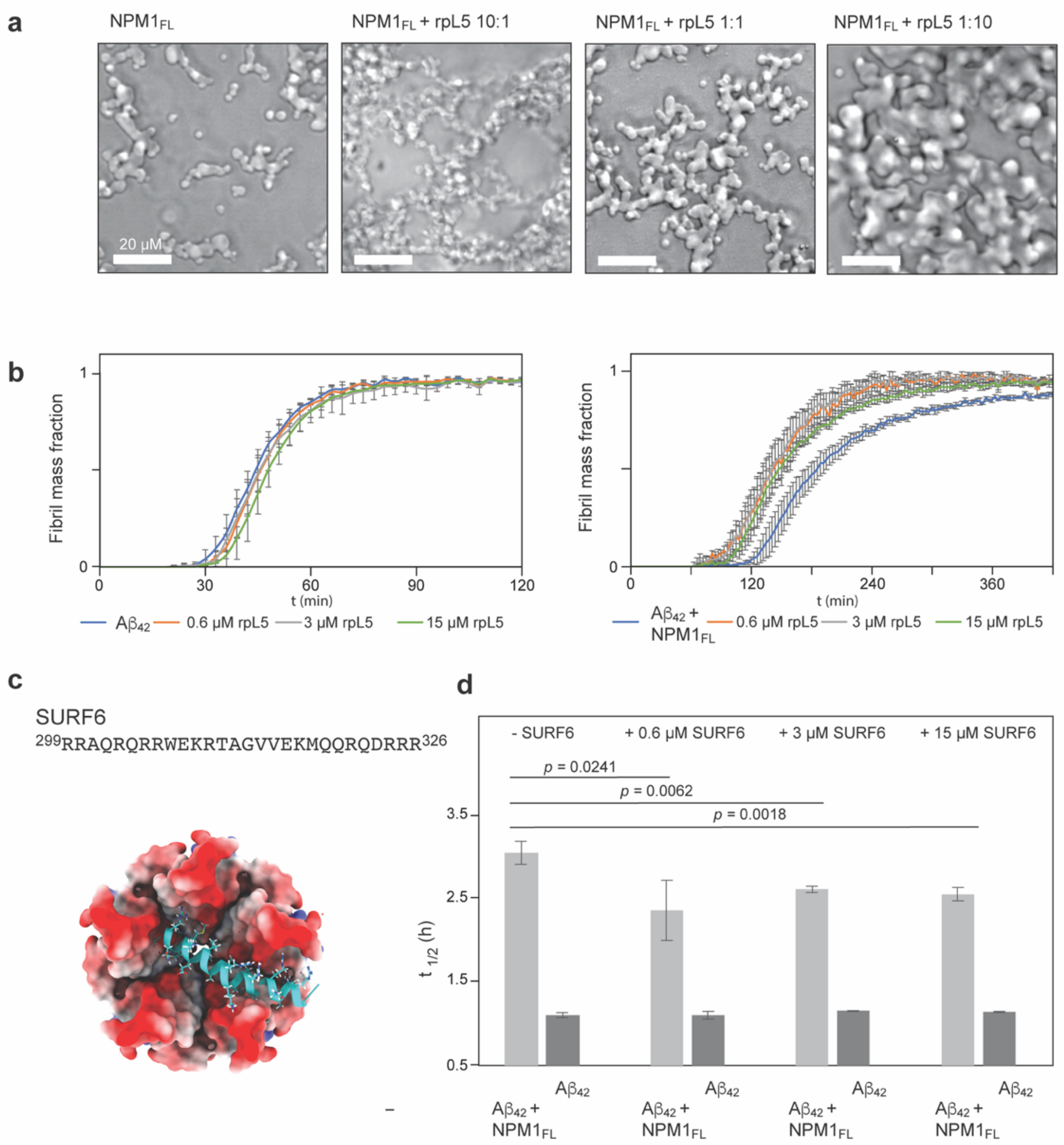


**Figure S7. AF predictions and Aβ$_{42}$ aggregation assays of NPM1 bound to rpL5$_{22\text{-}37}$ and SURF6$_{299\text{-}326}$.** (a) Light microscopy of FL NPM1 after incubation for 60 min alone or in the presence of rpL5$_{22\text{-}37}$ at ratios of 10:1, 1:1, or 1:10 shows an increase in number and size of NPM1 assemblies. (b) Left: ThT fluorescence curves for Aβ$_{42}$ alone as well as in the presence of rpL5$_{22\text{-}37}$ show no significant effect on fibril formation. Right: ThT fluorescence curves for Aβ$_{42}$ in the presence of FL NPM1 show reduced chaperone activity in the presence of rpL5$_{22\text{-}37}$. (c) AF prediction for the complex between NTD and SURF6$_{299\text{-}326}$ indicate binding of the SURF6 peptide to the acidic side of the NPM1 pentamer. The NTD pentamer is rendered as electrostatic surface, the SURF6 peptide is shown as ribbon in turquoise with basic residues as sticks. (d) T$_{1/2}$ of Aβ$_{42}$ aggregation curves reveal that SURF6$_{299\text{-}326}$ has no pronounced effect on Aβ$_{42}$ aggregation alone, but significantly reduces the chaperone-like activity of FL NPM1. Error bars indicate the standard deviation of n=4 repeats. Significance was calculated using Student's T-Test for paired samples with equal variance.

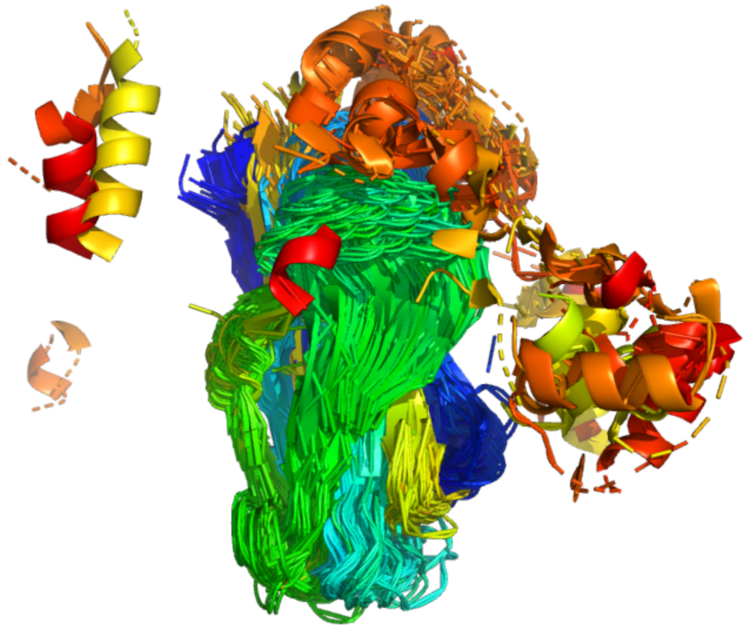

**Figure S8. AF models for 29 homologs of NPM1 with identical domain architectures.** Models are colored from N- to C-terminus and only non-disordered residues of the CTDs that are within a 4Å distance cut-off from the NTD are shown.
